# Hypervariable DOM properties in coastal NW Mediterranean Sea -Evidences of strong human influences and potential consequences for the heterotrophic base of planktonic food webs

**DOI:** 10.1101/2024.03.20.585671

**Authors:** Clara Dignan, Véronique Lenoble, Nicole Garcia, Benjamin Oursel, Ana-Marija Cindrić, Benjamin Misson

## Abstract

Marine DOM plays a key role in the carbon cycle. Though there is growing interest for its chemical and ecological properties, its variability in the very heterogeneous coastal environment is poorly documented. In this study, we assessed the spatial and temporal variations in marine coastal DOM chemical properties and its potential to support the growth of the heterotrophic basis of the planktonic food-web. For that, we considered two NW Mediterranean coastal sampling stations under contrasted continental and human influences. From January to July 2022, DOC concentration demonstrated a progressive increasing trend in addition to marked spatial differences. DOM composition presented more variations in time than in space. These variations appeared related to a combination of direct freshwater inputs tracked by salinity variations, direct human contaminations tracked by concentrations in Cu and Pb, and phytoplanktonic production. An experimental approach was also used at each sampling date to evaluate DOM ability to support the growth of heterotrophic prokaryotes. Significantly higher growth was observed with DOM from a site under higher continental and human influences. Water temperature exerted a higher control on growth than DOM properties. Correlation analysis with DOM sources suggested the uncoupling of phytoplanktonic production and growth of heterotrophic prokaryotes, which appeared better supported by human contaminations and, to a lesser extent, freshwater inputs. Sediment resuspension in harbors and antifouling paints could represent two important sources of bioavailable resources, favoring fast heterotrophic growth and higher net production, respectively.

## 1 Introduction

Marine DOM is one of the largest carbon pools on the planet (Hedges et al., 1992; Hansell et al., 2009) and an important and influential component of global carbon cycle models (Farrington, 1992). In the last decades, its spatial and temporal variability across ocean basins and its importance in the biological carbon pump were largely described (e.g. Hansell et al., 2009, Santinelli 2015). Its dynamics at various time scales is far from being fully understood. An ecological perspective linking production and consumption processes is emerging, although the role of key environmental drivers is debated (Shen and Benner, 2020, 2022; Dittmar et al., 2021; Lennartz and Dittmar, 2022). Through their major contribution to global ocean primary production (Gattuso et al., 1998), highly-dynamic marine coastal areas appear as hot spot of DOM production and processing. Along the coast, DOM originates from different sources. In agreement with the high primary production, an important part of coastal marine DOM is autochthonous, mainly resulting from phytoplanktonic activity. It varies along the year, in particular between the winter and summer periods because of ecological successions (Sommer et al., 2012; Romagnan et al., 2015; Yucel, 2018). Then, unlike offshore environments, coastal areas receive significant DOM inputs from allochthonous sources. Brought to the coastal environment by the drainage of the watershed, terrestrial DOM resulting from terrestrial biological production and its decomposition can feed coastal waters (Jonsson et al., 2017; Retelletti Brogi et al., 2021). Due to increasing demographic pressure, continental and marine human activities threaten marine coastal ecosystems (Halpern et al., 2008; The MerMex Group et al., 2011). This situation is particularly relevant around the Mediterranean Sea, given the human population constantly-increasing growth and footprint on its shoreline (UNEP/MAP and Plan Bleu, 2020). This represents a third source of DOM, non-natural, to the coastal environment: anthropogenic DOM, defined as the mixture of organic substances produced by human activities, brought by rivers and urban runoff, deposited from the atmosphere or directly discharged into coastal waters (Vila-Costa et al., 2020). With this diversity of DOM sources, the high spatial and temporal heterogeneity of coastal environments is now challenging our ability to precisely depict consequences of human influences from natural ones. Increasing scientific observations focused on DOM appears as a first step in the long way necessary to identify and then manage the most influent human activities.

Once produced or brought to the ocean, DOM fate is under the control of various factors among which heterotrophic prokaryotes (HP) are key players. Being the main DOM consumers, able to transform labile compounds into more recalcitrant ones, HP control the microbial carbon pump (Jiao et al., 2010) and reinforce the basis of planktonic food webs through heterotrophic assimilation. This heterotrophic base sustains higher trophic levels through the microbial loop (Azam et al., 1983).HP contribution to DOM fate is actually a part of a tryptic relationship with DOM properties and environmental conditions, both influencing HP metabolic activity and growth. Indeed, DOM intrinsic properties are known to influence its processing by HP communities (Amon and Benner, 2001; Shen and Benner, 2020). Abiotic conditions, such as temperature that can control HP metabolic activity (Pomeroy and Wiebe, 2001) or inorganic nutrients availability (Thingstad et al., 1997), are also known for their influence on DOM processing by HP. Then, three main biotic characteristics of the marine environment are also contributing. First, HP development is under strong top-down pressure by grazers and viruses in the marine environment, limiting DOM processing rates (e.g. Liu et al., 2014; Silva et al., 2019). Secondly, competition with phytoplankton for inorganic nutrients can reduce the ability of HP to efficiently use DOM (Thingstad et al., 2005). Last but not least, DOM reworking by HP communities themselves tends to lower DOM lability and furthers contributes to limit DOM processing by HP (Mentges et al., 2019 ; 2020). Exposed to a diversity of processes, these numerous controls of DOM fate can result in major changes in biogeochemical functioning. In the coastal environment, terrestrial discharge is for example known for its ability to shift the basis of planktonic food webs from a mainly autotrophic to a mainly heterotrophic metabolism (Wikner et al., 2012; Jonsson et al., 2017, Barrera-Alba et al., 2019; Navarro et al., 2023). Numerous human activities on the coasts, especially concentrated in large, urbanized bays, could also influence biogeochemical fluxes trough imbalances in DOM properties. Here again, an important observational effort is now needed understand and model coastal DOM dynamics in this highly variable context for a more sustainable management of human activities.

In this context, this study aimed at documenting spatiotemporal variability in DOM properties in a Mediterranean coastal area, to compare contributions of phytoplanktonic, continental and human influences and to further provide pieces of evidence of ecological consequences. For that, we combined (i) in situ sampling for the characterization of DOM chemical properties and tracers of autochthonous, continental and anthropogenic sources and (ii) experimental assessment of DOM potential to support HP growth under low top-down pressure. We observed a very large variability in DOM chemical properties with unexpected short-term and spatial changes that override seasonal trends. DOM ability to support the heterotrophic basis of the food web appeared largely controlled by continental and human influences, and strongly modulated by water temperature. This work suggests that human activities and constructions in harbors have the potential to strengthen the heterotrophic basis of the planktonic food web.

## 2 Material and methods

### 2.1 Study area and sample collection

The first coastal sampling station was located in Niel Bay on the Giens peninsula (“O”, 43.035325; 6.127527), within Port-Cros National park, located along the French coasts in the North-Western Mediterranean, in an area open to the sea and regularly stirred by the currents.

This station is exposed to very limited human pressure and its location in a rocky peninsula reduces strongly any continental influence. To maximise the range of potential variability in DOM properties, a second sampling station was chosen in Toulon bay, a site exposed to high levels of anthropogenic activities (1^st^ French naval base, ferry transport, civil harbours, industry, sewage and aquaculture). Human activities in this area have generated multiple chemical contaminations in the water as well as in the sediment (Tessier et al., 2011; Coclet et al., 2018, Layglon et al., 2020) and influence biological functioning (Coclet et al., 2018, 2019, 2020; Paix et al., 2021). This bay is also under the influence of freshwater discharge, mainly through 2 river outlets (Las and Eygoutier river; Durrieu et al., 2023; Nicolau et al., 2012). In this wide bay, the sampling station was more specifically located in the civil harbour of Toulon (“H”, 43.118460; 5.934092), which represents typical civil harbours of the French Mediterranean coast (in the centre of a city, landlocked, with very limited tides, hosting leisure boats as well as artisanal fishing activity). Both sites being located within a restricted geographic area (∼ 18 km), they were exposed to similar climatic conditions.

These two stations were sampled every two to three weeks, from January 2022 to July 2022. Standard oceanographic properties including water temperature (°C), salinity and chlorophyll a (µg.L^-1^) were measured using a multiparameter probe (Hydrolab® DS5X). For the different chemical analyses, surface seawater (1 m depth) was sampled at each site with previously washed and conditioned material (see Supplementary Material). One liter of seawater was sampled using a Van-Dorn bottle and transferred in a FEP bottle for chemical characterization. Similarly, 15L of surface seawater were collected at each site and transferred in a 20L LDPE bottle for incubation experiments. In order to avoid any major changes in the DOM properties and the prokaryotic communities, the samples were brought back in the lab within 2 hours and processed immediately. To characterize dissolved substances, subsambles were taken in the 1L FEP bottle as follows: 24 mL were filtered through 0.2 µm polyesthersulfone (PES) syringe filters, collected in preconditioned glass tubes, acidified (10% v/v HCl, analytical grade, Fisher Scientific) and stored at 4°C waiting for dissolved organic carbon (DOC) concentration measurement. Following the same filtration protocol, 24 mL of filtered water were stored in preconditioned glass tubes at 4°C for further fluorescent DOM (fDOM) evaluation and 50 mL were stored at -20°C in centrifuge tubes (Falcon) for nitrate, nitrite, dissolved organic nitrogen (DON), phosphate and dissolved organic phosphorus (DOP) concentration measurements.

Ten mL of filtered water were also collected in Trace Metal Grade centrifugation tubes (Falcon®), acidified (0.2% v/v HNO_3_ sp, analytical grade, Fisher Scientific) and stored at 25°C waiting for copper (Cu) and lead (Pb) concentration measurement by HR-ICP-MS.

#### Experimental design

In order to set an experimental approach representative from in situ conditions, we incubated natural microbial communities in natural dissolved substances pools. For that purpose, at each sampling date, heterotrophic microbial communities and dissolved substances from each site were isolated by filtration. Seven hundred millilitres of seawater were filtered through a pre- conditioned PES filter (Whatman, 0.2 µm, 47 mm) to recover pools of dissolved substances. In parallel, 100 mL of seawater were filtered through a pre-conditioned GF/F (Whatman, 0.7 µm, 47 mm) to isolate the free heteretrophic prokaryotic (HP) community from each sample. The filtrates were stored at site temperature until the experimental mixtures were processed.

In the lab, the experiments consisted in incubating each pool of dissolved substances to the HP community of the same sampling site. In addition, each dissolved pool was also incubated with the HP community of the other site to evaluate the consistency of DOM potential to support HP growth. The experimental conditions were named “HinH” for the harbour community in its own dissolved substances, “OinO” for the open area community in its own dissolved substances, “HinO” for the harbour community in open area dissolved susbtances and “OinH” for the open area community in harbour dissolved substances This experimental approach developed within a previous work (Dignan et al., submitted) was hereby adapted to take into account the influence of *in situ* temperature and potentially delayed microbial growth by winter temperature. Thus, incubations lasted 72 h and were performed in the dark at the *in situ* temperature recorded during the sampling (or at the average temperature in case of difference between the two sampling stations). Incubations resulted in 4 triplicated experimental conditions at each sampling date. For each replicate, 54 mL of sterile seawater (< 0.2 µm) containing the dissolved substances was inoculated with 6 mL of seawater containing the HP community (< 0.7 µm) in a preconditioned 60 mL FEP bottle.

Subsamples (1 mL) were taken after 24, 48 and 72 h of incubation, fixed with glutaraldehyde (0.25% final concentration) and stored at -80°C prior to HP enumeration by flow cytometry. Differences in net growth and in particular the maximum heterotrophic prokaryote abundance (mHPA) reached during incubation were used as proxies to compare DOM potential to support HP growth.

### 2.2 Chemical analyses for field samples

To evaluate the quantitative properties of DOM present at each site, DOC concentrations were determined by high temperature catalytic oxidation using a Shimadzu TOC-VCSH carbon analyzer with an accuracy of 0.1 mgC.L^-1^. DOC analysis were validated by comparison with the DOC consensus reference material (SUPER-05) (Hansell, 2005; Louis et al., 2009). Nutrients (nitrate NO_3-_ and phosphate PO_43-_) were measured by colorimetric methods using an automated Technicon Autoanalyser III (Treguer and Le Corre, 1975) and total matter concentrations (total nitrogen TN and total phosphorus TP) were determined using the wet-oxidation technique (Raimbault et al., 1999). Dissolved organic nitrogen (DON) and phosphate (DOP) concentrations were deduced by subtracting nitrate concentration from total nitrogen concentration and phosphate concentration from total phosphorus concentration, respectively.

DOM quality was estimated using the composition of DOM fluorescent fraction. Three-dimensional excitation-emission matrices (EEM) of each sub-sample were recorded using a Fluoromax+ spectrofluorimeter (Horiba) using a 1 x 1 cm quartz cell. EEMs are used to distinguish fluorescence signals due to various groups of chromophores. The excitation wavelength varied between 220 and 450 nm in 5 nm increments, while the emission was measured between 220 and 600 nm. By using the drEEM toolbox (Murphy et al., 2013), the EEMs were corrected from the Milli-Q water EEM measured under the same conditions. EEMs were elaborated in order to remove and interpolate the Rayleigh and Raman scatter peaks. EEMs were normalized to the Raman signal of water, dividing the fluorescence by the integrated Raman band of Milli-Q water (λex = 350 nm, λem = 371–428 nm), measured on the same day of analysis. Thus, fluorescence intensity is expressed in water-equivalent Raman units (R.U.) (Lawaetz and Stedmon, 2009). PARAFAC analysis was performed on all EEMs collected during time tracking in MATLAB R2018a (Mathworks, Natick, MA). PARAFAC analysis was performed using two to seven component models with non-negativity constraints. The final four-component model (Fig. S.I. 1) was chosen based on residual analysis, split-half analysis, and visual inspection (Stedmon and Bro, 2008).

Two trace metals considered as proxies of different human influences in Toulon Bay were considered for this study, Copper (Cu) and lead (Pb), as Toulon Bay was demonstrated to present an important metallic contamination (Tessier et al., 2011). Cu contamination is linked to the use of antifouling paints (Lageström et al., 2020). The long residence time of the waters within the bay coupled to the absence of tides of the Mediterranean Sea leads to an important contamination of the waters, fluctuating with the seasons and the hydrodynamics of the area (Mazoyer et al., 2020). As a consequence, Cu appears strongly enriched in harbours of the bay (Coclet et al., 2018). Hotspots of sediments PB contamination were highlighted near Toulon, resulting from Second World War boats scuttling (Dang et al., 2015). Through waves, boat traffic or dredging, sediments are remobilised in the water, leading to some metals’ desorption, especially lead (Dang et al., 2020; Layglon et al., 2020). The concentrations of dissolved Cu and Pb were measured by High Resolution Inductively Coupled Plasma Mass Spectrometer (Element 2, HR ICP-MS, Thermo), with a 10-fold dilution for seawater samples to reduce salt- matrix effect. All samples were spiked with an Internal Standard (Indium). A certified reference material (CASS-5, Nearshore seawater reference material for metals, National Research Council Canada) was used as a quality control of HR ICP-MS measurements. Two separate CASS-5 control samples were measured after every 10–15 samples. Matrix matching calibration (in 10-fold diluted CASS-5 sample) was used for concentration quantification. Determined concentrations of metals in CASS-5 sample were within 10% of the certified reference.

### 2.3 Heterotrophic prokaryotes abundances (HPA) during incubations

After thawing, samples were stained with SYBR Green (1X final concentration) for 15 minutes in the dark and HP were counted with an Accuri C6 flow cytometer (BD Biosciences). Fifty microliters were run at a flow rate of 35 µL.min^-1^. Non-fluorescent polystyrene microspheres (Flow Cytometry Size Calibration Kit, Thermo Fisher Scientific) were used as a size standard. Particles considered as HP were smaller than 2.0 µm, exhibited low complexity (low SSC), emitted green fluorescence and no red fluorescence (Grégori et al., 2001). Data were acquired using BD Accuri CFlow Plus software and HPA expressed as cells per milliliter (cell.mL^-1^). Maximal HP abundances (mHPA), always recorded after 72h, were used for interpretations.

### 2.4 Statistical analyses

All statistical analyses were performed with RStudio 2022.7.0.548 software (RStudio Team, 2022) under R 4.2.0 environment. The variability of DOM properties according to sampling stations was represented through PCA with *stats* and *ggplot2* packages. Significant contribution of the represented variables was verified with the *envfit* function of the *vegan* package.

After verifying the normal distribution and homoscedasticity of the variances with Shapiro and Levene tests, respectively, analyses of variance (ANOVA) were used to compare the mHPA obtained under each experimental condition at every sampling date using rstatix package. If the ANOVA detected a significant difference (p-value < 0,05), a pairwise T test, with the Bonferroni method for the p-value correction, was then used to distinguish which samples were significantly different from the others (p-value < 0,05). Wilcoxon tests were used to compare the average values of environmental variables between the two sites over the entire monitoring period.

To investigate the relationships between DOM properties and potential sources of variation, Spearman rank pairwise correlation tests were performed using the “vegan” package in R. Corresponding heatmaps were built using ggplot2. For specific correlation analysis between environmental variables and HP growth, we considered growth kinetics in addition to mHPA. For that, daily net growth were calculated by substracting HPA recorded at the beginning of the 24h time frame to HPA recorded at the end. Then, Spearman rank pairwise correlation tests were performed and represented through heatmaps, as described above.

## 3 Results

### 3.1 Spatial and temporal variability in DOM chemical properties and potential sources

Overall variability in DOM properties (DOC, DON, DOP and fluorescent components) at the two studied stations throughout the temporal sequence were visualized by a Principal Component Analysis (Fig. 1). The two-dimension representation covered ∼70% of the observed variations. Such representation allows to highlight an important temporal variability, which is accentuated at station H, the temporal sequence spreading far more than those from station O. There is no seasonal trend, whatever the considered station, pointing to occasional variations. Such variability appeared to be mainly driven by DOC, DON and three fluorescent components.

**Figure 1.**
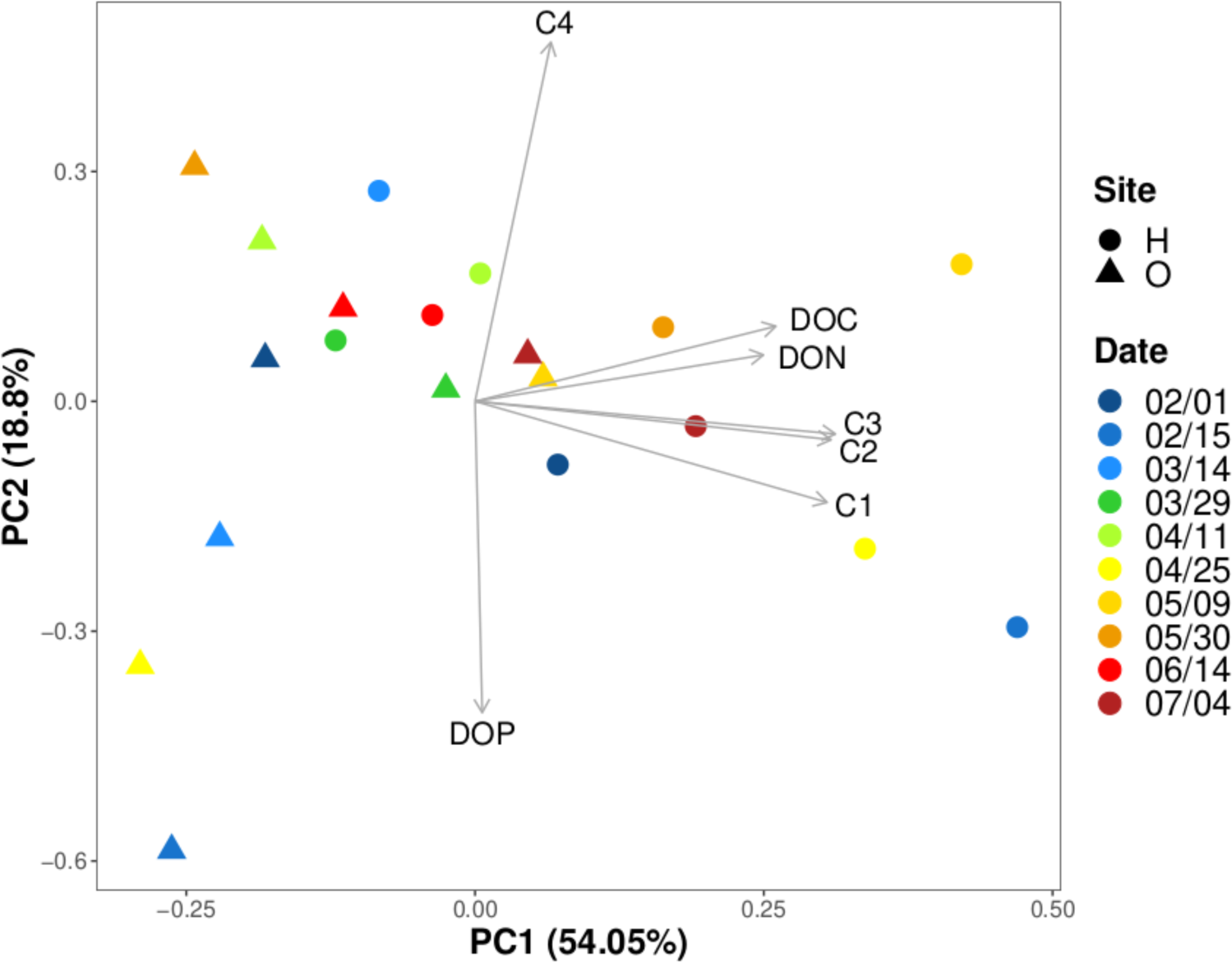
Principal component analysis of the variation of DOM properties through time and space. H: harbour; O: open area. Indicated sampling dates correspond to 2022.

During the monitoring, DOC showed significant variations at both stations (from ∼ 60 to ∼85 µM at station O, and from ∼70 to 105 µM at station H; Fig. S.I. 2). Such values are in the range of the reported values for this long-time studied area (Dang et al., 2018; Coclet et al., 2019; Layglon et al., 2022; Durrieu et al., 2023). The average DOC concentrations increased throughout the temporal sequence. While average DOC concentration appeared significantly higher at H (p-value < 0.05), temporal variations within a given station were of similar extent to inter-stations variations (∼ 15 µM). DON and DOP concentrations varied between ∼ 4 to 15 µM and ∼ 0.1 and 0.3 µM, respectively. These ranges are in line with previous observations in another NW Mediterranean coastal site located close to an urbanized bay (Céa et al., 2015). No clear temporal trend could be observed (Fig. S.I. 2), and the temporal punctual variations within one station were more important than inter-stations variations. Differences in average concentrations in DON and DOP between H and O were not statistically significant (p-value > 0.05).

The PARAFAC analysis performed on the EEMs allowed to identify four fluorescent components (Fig. S.I. 1). From the literature and the OpenFluor database, component 1 corresponded to marine humic-like organic matter (C1_MarHum-like_), component 2 to tryptophan-like (C2_Trp-like_), component 3 to terrestrial fulvic-like (C3_TerFul-like_) and component 4 to tyrosine-like (C4_Tyr-like_). At station O (Fig. S.I. 2), the fluorescence intensity of the different compounds showed very limited variation. At station H, C1_MarHum-like_, C2_Trp-like_ and C3_TerFul-like_ were significantly enriched when compared to station O (p < 0.01). Components C2_Trp-like_ and C3_TerFul-like_ were highly variable (Fig. S.I. 2), which is in line with the PCA results.

These numerous spatial and temporal variations in DOM chemical properties could reflect different processes and sources. First, significantly higher concentrations in chlorophyll a were observed at H (p-value < 0.001), which is in agreement with previous observation of phytoplanktonic diversity suggesting higher production in the most enclosed parts of Toulon Bay (Coclet et al., 2018; Delpy et al., 2018). Then, other sources could explain these higher concentrations, like the input of freshwaters or the concomitant input of organic matter and contaminants related to human activities. Salinity was significantly lower and trace metallic contamination was significantly higher at H (p-value < 0.05). Most of salinity variations could be related to the difference in land influence between the two sampling stations, illustrated by the presence of two rivers (Las and Eygoutier) in Toulon Bay. They are characterized by a usually small discharge, except during the typical Mediterranean violent but short storms (Durrieu et al, 2023; Nicolau et al., 2012). Station H being located within Toulon harbor in the Nort-Western part of Toulon Bay and station O appearing far more remoteness, spatial differences in trace metallic contamination are in good agreement with previous observations in the area (Coclet et al., 2018, 2019, Layglon et al., 2020). Concerning temporal variability, salinity, Chl a and trace metals did not fluctuate much at station O when strong variations were recorded at station H (Fig. S.I. 3). As previously explained, station O is prone to very limited continental and human influences, which explains these small fluctuations. On the other hand, station H is exposed to both higher continental and human influences, as well as located in an area showing high variability in hydrodynamics (Dufresne et al., 2014; Mazoyer et al., 2020).

As all these processes and sources could influence DOM chemical properties, a correlation analysis was performed (Fig. 2). Looking at DOM quantitative properties, DON and DOP did not present any significant correlation with any of the studied source tracers. Salinity was negatively correlated with DOC, suggesting DOC-rich freshwater inputs. Terrestrial discharges through freshwater often represent an important source of DOC to the coastal sea, as observed elsewhere in the oligotrophic Mediterranean Sea (e.g. Navarro et al., 2023). Then the positive relationship between Cu and DOC let us hypothesize that they have a common source: the antifouling paints. As a matter of fact, the new generation of antifouling paints are designed to be self-polishing (Watermann & Eklund, 2019), which means that they are progressively degrading through hydrolysis, releasing a whole set of compounds in the water, including some organic compounds from the polymeric matrix (Yebra et al., 2005). Surprisingly, DOC concentration was not correlated to chlorophyll concentration, an observation that sets the difference with other Mediterranean contexts (Avril, 2002; Santinelli et al., 2015, 2020).

**Figure 2.**
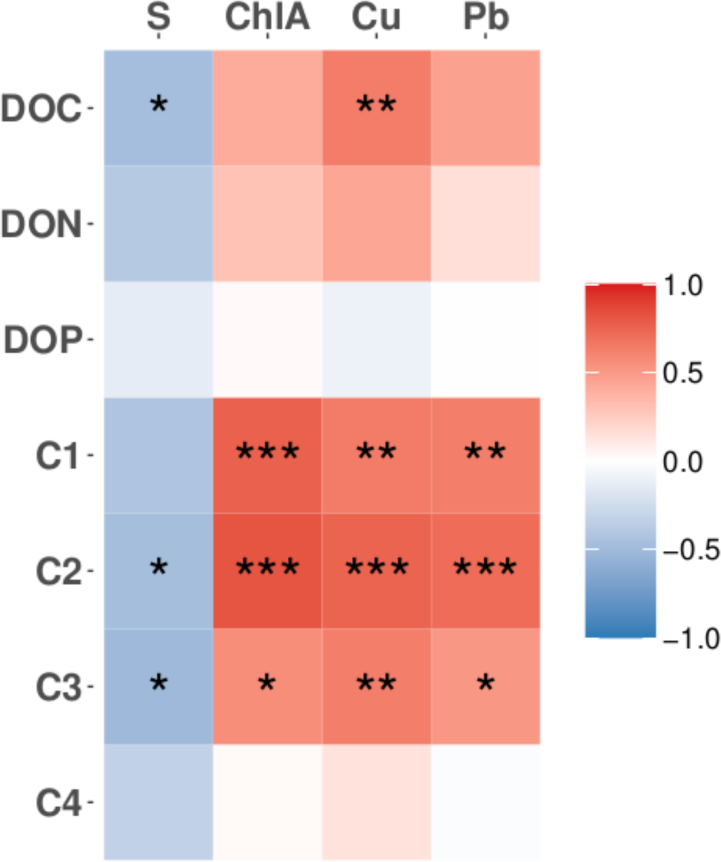
Correlation analysis between variations in DOM chemical properties and the considered sources. C1 to C4 are representing the fluorescent components identified with PARAFAC: marine humic-like, tryptophan-like, terrestrial fulvic-like and tyrosine-like, respectively. The color scale corresponds to Spearman’s correlation coefficients. 1, 2 and 3 stars represent a p-value < 0.05, < 0.01 and < 0.001, respectively.

The composition of DOM, expressed through fluorescent components C1_MarHum-like_ and C2_Trp-like_, was highly linked with chlorophyll (Fig. 2). This strongly suggest the influence of fresh phytoplanktonic production of DOM and heterotrophic reworking on its chemical quality. Indeed, protein-like fluorescent component (such as C2_Trp-like_ in this study) is known to be produced and exudated during marine phytoplankton exponential growth (Bachi et al., 2023). Moreover, marine humic-like fluorescence (C1_MarHum-like_) is often associated with autochthonous fluorescent DOM and its presence has historically been linked to elevated biological activity in the water column (Nieto-Cid et al., 2005; Stedmon and Markager, 2005; Romera-Castillo et al., 2010). Then, negative correlations between fluorescence intensities and salinity suggest an indirect contribution of freshwater inputs that could stimulate phytoplanktonic production. The correlation between salinity and fluorescent component C3_TerFul-like_ support the hypothesis of influent continental inputs of DOM. Lastly, DOM composition appeared related to the studied metals. These correlations being positive, any toxic influence can be neglected but such covariations support common sources. This strongly support the hypothesis of a strong impact of antifouling paints and sediment remobilization on DOM chemical properties.

In the studied area, DOM is thus presenting a huge variability, partly linked to the temporal sequence. It has to be underlined that the temporal qualitative variations in a considered station largely overpass the inter-station variations. This statement is exacerbated at station H, a more land-connected area exposed to numerous anthropogenic activities. DOM quantity and properties are deeply linked with these covarying potential sources of direct and indirect influences.

### 3.2 Stimulation potential of the heterotrophic basis of the planktonic foodweb

During incubation experiments, HPA increased in all conditions. Several growth phases were observable: an exponential growth phase before growth slowed down when the community reached the carrying capacity of the system (Fig. S.I. 4). However, from January 18^th^ to March 29^th^ no plateau was observed, suggesting that the carrying capacity might not have been reached after three days of incubation, the communities being still in the exponential growth phase. In comparison of the potential dissolved substances to sustain heterotrophic growth, the maximum Heterotrophic Procaryotic Abundance (mHPA) reached in each incubation was used. It was systematically reached after 72h of incubation.

The use of dissolved substances from H resulted in a high temporal variability (HinH: 1.38 ± 0.88x10^6^ cell.mL^-1^, OinH: 1.09 ± 0.90x10^6^ cell.mL^-1^) and promoted the highest mHPA (Fig. 3). With dissolved substances from O, variations were smaller though still important (HinO: 5.49 ± 0.25x10^5^ cell.mL^-1^, OinO: 3.14 ± 0.22x10^5^ cell.mL^-1^). No clear temporal trend could be depicted, and several punctual important variations were observed. Average mHPA appeared significantly lower with dissolved substances from O than with those from H (p-value < 10^-4^). This observation agrees with previous observations in the same geographic area (Dignan et al., submitted). Differences in growth stimulation between the two pools of dissolved substances were well supported whatever HP origin from April to July. However, the influence of HP origin was predominant from January to March. While it has been proposed in the literature that a given DOM pool could be more available to some HP communities than others (e.g. Carlson et al., 2004), longer incubations allowing to reach the carrying capacities would have been necessary to validate this conclusion. Nevertheless, the observed delayed growth at that period suggests at least a higher reactivity of HP from H. More work is now needed to explore this trend associated to winter and rather cold waters.

**Figure 3.**
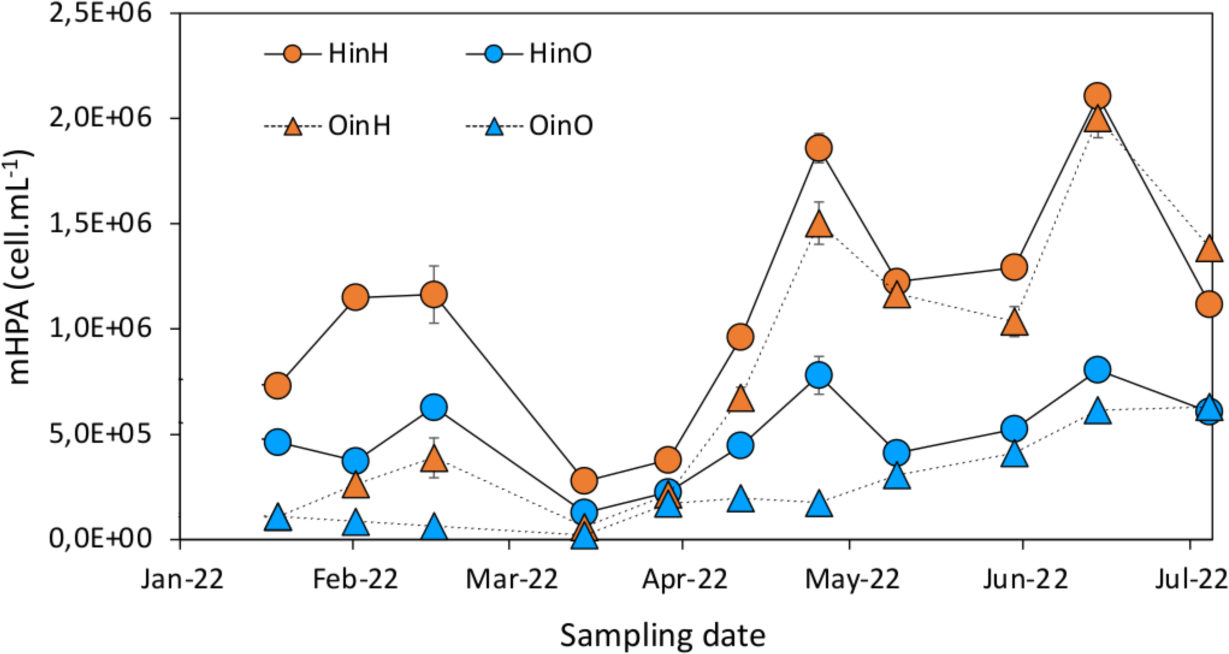
**Maximum heterotrophic prokaryotes abundance (mHPA) as a function of DOM origin for each campaign**. HP origin is specified by point shapes, circles: Harbour (H); triangles: Open area (O). DOM origin is specified by point colours, orange: Harbour (H); blue: Open area (O). Error bars represent the standard deviation between experimental triplicates.

Utilization of marine DOM by HP for biomass and energy production is known to be determined by DOM quantitative and qualitative properties (Del Giorgio and Cole, 1998; Amon and Benner, 2001; Shen and Benner, 2020), but inorganic nutrients supply can also play a key role when DOM composition does not meet HP stoichiometric requirements (Thingstad et al. 1997 AME). Moreover, through its influence on microbial metabolism, water temperature is known to regulate HP growth rates (Pomeroy and Wiebe, 2001), especially when nutrient supplies are not limiting (Thingstad and Asknes, 2019). To better evidence the abiotic variability that could explain the observed variations in HP growth according to time and space, we also considered water temperature, as well as nitrate and phosphate concentrations in addition to environmental variables presented above (Fig. S.I.5). Seawater temperature gradually increased over the studied period, rising from ∼ 13 °C in January to ∼ 25 °C in July. It presented similar temporal variations at both stations, H being slightly colder in winter and warmer in summer. Phosphate concentrations measured at H (0.06 ± 0.11 µM) were similar to previous measurements in another NW Mediterranean site (Céa et al., 2015). They were significantly lower at O (p-value < 0.01) and very low compared to previous observations (0.01 ± 0.01 µM; Céa et al., 2015). While it appeared highly stable at O, important punctual increases were observed at H, reaching concentrations up to 20 times higher than at O (Fig. S.I.5). During the studied period, nitrate concentrations remained low and stable at O (0.29 ± 0.28 µM), while they were significantly higher (p-value < 10^-4^), hugely variable (6.73 ± 8.10 µM, Fig. S.I.5) and elevated at H when compared to another French Mediterranean coastal station under the influence of an urbanized bay (Céa et al., 2015).

Correlation analyses between HP growth metrics and environmental variables in the whole dataset highlighted numerous highly significant correlations (Fig 4). The correlation with water temperature is in good agreement with its well-known effect on microbial metabolism and growth (Pomeroy and Wiebe, 2001). Considering DOM properties and inorganic N and P supplies, DOC concentration appeared as the most influent factor for growth kinetics and total net growth (Fig 4), growth being faster and more important when DOC concentration was higher. This observation is supporting the debated role of DOC concentration on microbial behaviours (e.g. Shen and Benner, 2020, 2022; Lennartz and Dittmar, 2022). Then, qualitative properties of DOM also appeared influent, especially stimulating mHPA, when C1_MarHum-like_, C2_Trp-like_ and C3_TerFul-like_ fluorescence and DON concentration were higher (Fig 4). The influence of nitrate concentration appeared similar to these DOM properties. These observations are in line with the influence of DOM qualitative properties on HP growth (Del Giorgio and Cole, 1998; Amon and Benner, 2001; Shen and Benner, 2020). Phosphorus content of DOM did not present any correlation with HP growth in our study. Similarly, inorganic phosphorus presented very weak correlations with growth (Fig 4). Although the weak apparent influence of phosphorus could appear quite surprising when considering the well-established phosphorus limitation of HP growth in the Mediterranean Sea (e.g. Céa et al., 2015, van Wambeke et al., 2002), this observation suggests that the proximity of the shoreline in both studied sites ensured sufficient phosphorus loading. Finally, correlations between HP growth metrics and nitrate, DON and FDOM were weaker and no longer significant when considering a single sampling station (data not shown). Thus, these substances would more be responsible of spatial variations in the potential of dissolved pools to sustain the heterotrophic growth at the basis of the planktonic foodweb than the temporal one. On the other hand, temperature and DOC concentration still presented significant and strong correlation, whatever the station, suggesting their combined influence on temporal variabilities in HP growth sustainment.

**Figure 4.**
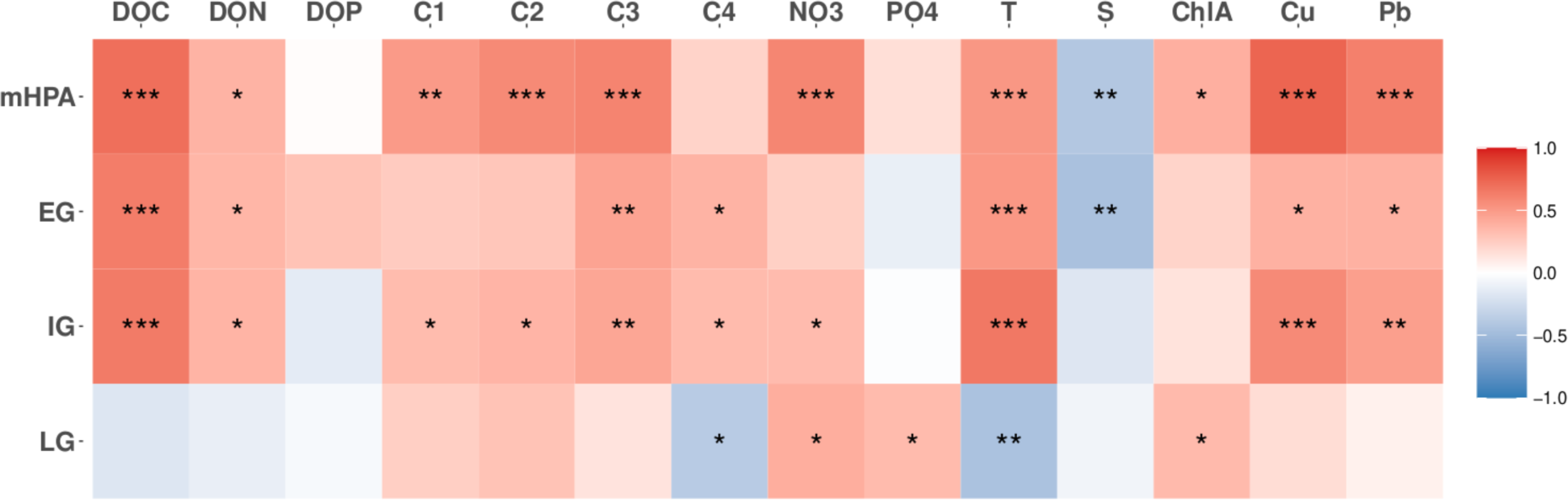
Correlation analysis between HP growth, DOM properties, other constraining factors and the considered tracers of potential sources of dissolved resources. mHPA: maximal heterotrophic prokaryotes abundance, EG: early growth (between 0 and 24h), IG: intermediate growth (between 24 and 48h), LG: late growth (between 48 and 72h). The color scale corresponds to Spearman correlation coefficients. 1, 2 and 3 stars represent a p-value < 0.05, < 0.01 and < 0.001, respectively.

Correlations with tracers of potential sources of dissolved sources highlighted interesting patterns. Indeed, chlorophyll a appeared very poorly correlated to HP growth metrics whereas salinity was very significantly and negatively correlated to early and total net growth (Fig 4). This observation suggest that HP growth was more stimulated by freshwater inputs than by variations in phytoplanktonic production, even though salinity variations were small. Similarly, Cu and Pb showed far stronger and more significant correlations with early and total net growth than chlorophyll a (Fig 4). Such correlations being positive, they demonstrate the absence of toxicity of these metals in our study and suggest that the heterotrophic base of the planktonic food web in the considered marine area could largely be sustained by human activities. While it is generally considered that phytoplanktonic production is the main source of organic substrates for HP growth in oligotrophic marine environments, shifts in the autotrophic/heterotrophic balance have been observed in environments under strong allochthonous inputs such as estuaries (Barrera-Alba et al., 2009; Jonsson et al., 2017; Wikner & Andersson, 2012). Our results strongly support the idea of such shifts in the studied area, even in the absence of major riverine inputs, and points out the strong influence of human activities on biogeochemical functioning at the basis of the planktonic foodweb. Such impact could be linked to direct inputs of dissolved substances but also to increased water residence time because of human constructions, and subsequent accumulation favouring eutrophication. Indeed, in the same area, an important increase of water residence time has been demonstrated at rather large scale under the influence of the main seawall of Toulon Bay (Dufresne et al., 2014), an influence that is probably even more important in enclosed docks such as Toulon civil harbour.

To get more insights into human influences, another correlation analysis was performed focusing on the results obtained with water from H, the harbour being exposed to multiple human influences (Fig. 5). This analysis confirmed the absence of significant correlation with chlorophyll a. Moreover, it did not show any correlation with salinity, suggesting that the above-mentioned significant correlations were mainly linked to differences among the chosen sampling stations. Interestingly, Cu and Pb demonstrated different significant and positive correlations with growth metrics, higher concentrations in Cu being associated to higher total net growth and higher Pb concentrations being associated to higher early growth. In Toulon Bay, the highest concentrations in dissolved Cu concentrations were observed in civil harbours (Coclet et al., 2018, 2019), suggesting antifouling paints as a major source for this contamination. Considering that, our study tends to demonstrate that the concentration of boats in civil harbours could additionally be responsible for organic contamination of the water with consequences for biological functioning. Concerning Pb, it was well demonstrated in the area by field work as well as lab experiments that sediment resuspension was a major source of dissolved Pb (Dang et al., 2015, 2020; Layglon et al., 2020). Our work then suggest that such resuspension influences dissolved nutritive substances in the water column and affect biogeochemical functioning at the basis of the planktonic foodweb.

**Figure 5.**
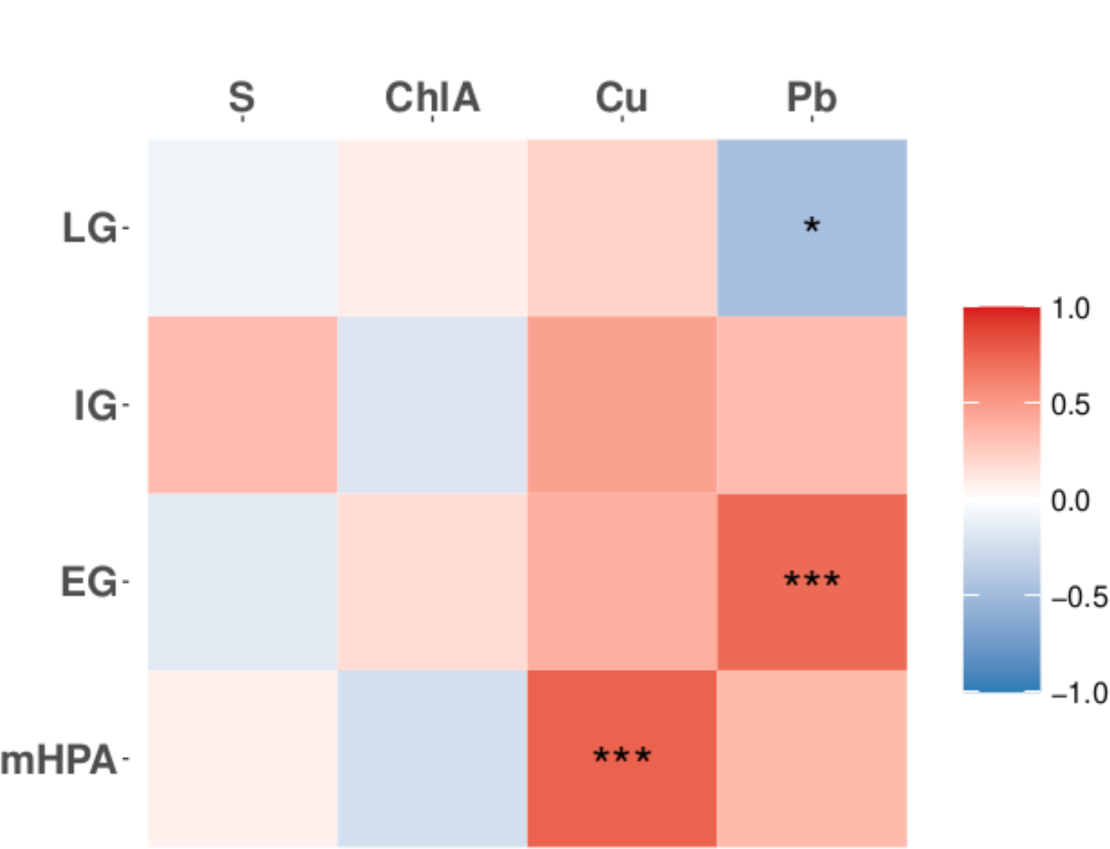
Correlation analysis between HP growth with water from Toulon harbour and considered tracers of potential sources of dissolved resources. mHPA: maximal heterotrophic prokaryotes abundance, EG: early growth (between 0 and 24h), IG: intermediate growth (between 24 and 48h), LG: late growth (between 48 and 72h). The color scale corresponds to Spearman correlation coefficients. 1, 2 and 3 stars represent a p-value < 0.05, < 0.01 and < 0.001, respectively.

## Conclusion

This study evidences a strong variability of DOM chemical properties in a coastal NW Mediterranean area. Important temporal changes were observed, with a seasonal pattern for DOC concentration and punctual changes in DOM qualitative properties. Such changes were amplified in a nutrient-rich harbour when compared to an oligotrophic open coastal site. Consequently, DOM ability to support heterotrophic growth at the basis of the planktonic food web was far higher when originating from the harbour, and more linked to quantitative DOM properties than to qualitative ones under our experimental conditions. Continental and anthropogenic sources appeared to be the main support of heterotrophic growth, which was additionally regulated by water temperature.

## Acknowledgements

This project was financially supported by Toulon University and the CARTT of the University Institute of Technology of Toulon and by the European Interreg Italy-France Maritime 2014- 2020 Project «GEREMIA» (Gestione dei reflui per il miglioramento delle acque portuali) and by the Regional Council of Provence Alpes Côte d’Azur (France). Clara Dignan received a Ph.D. fellowship from the Ministère de l’Enseignement Supérieur et de la Recherche and this paper is a part of her Ph.D.

