## Supplementary material for "Hypervariable DOM properties in coastal NW Mediterranean Sea -Evidences of strong human influences and potential consequences for the heterotrophic base of planktonic food webs"

### Pre-cleaning protocol

The water samples were taken with a horizontal van Dorn type water sampler (Wilco, model Beta) previously cleaned with acid (10% v/v HCl, analytical grade, Fisher Scientific) and thoroughly rinsed with Milli-Q water (18.2 MΩ, Millipore) in the laboratory. In the field, the sampler was rinsed with seawater from the site to be sampled before sampling.

Fifteen litres of seawater were stored in a twenty-litter bottle and one litter was stored in a fluorinated ethylene propylene (FEP) bottle. These containers were, in the laboratory, previously rinsed three times with Milli-Q water (18.2 MΩ, Millipore), sterilized with acid (10% v/v HCl, analytical grade, Fisher Scientific) for 24 hours under stirring, rinsed again three times with Milli-Q water (18.2 MΩ, Millipore) then filled with acid (0.1% v/v HCl, Trace Metal Grade, Fluka). On field, the FEP containers and bottles were rinsed three times with seawater from the site to be sampled.

All the FEP bottles used for incubation experiments followed the same conditioning protocol described above.

Dissolved substances and heterotrophic communities were isolated by filtration during the experiment. In order to avoid possible contamination of the carbon filtrates, the 0.2 µm polyesthersulfone filters (Whatman, 47 mm) were previously washed with 100 mL of acid (10% v/v HCl, analytical grade, Fisher Scientific), rinsed with 1L of Milli-Q water (18.2 MΩ, Millipore), then packaged with 150 mL of the sample to be filtered. The GF/F glass fiber filters were previously calcined (450°C, 6h), rinsed with 1L of Milli-Q water (18.2 MΩ, Millipore) then conditioned with 150 mL of the sample to be filtered.

The 24 mL glass tubes used for the DOC concentration and DOM fluorescence intensity analysis were previously rinsed three times with Milli-Q water (18.2 MΩ, Millipore), sterilized with acid (10% v/v HCl, analytical grade, Fisher Scientific) for 24 hours, rinsed again three times with Milli-Q water (18.2 MΩ, Millipore) then calcined (450°C, 6 hours). At the time of sampling, they were rinsed three times with the filtered sample before being filled.

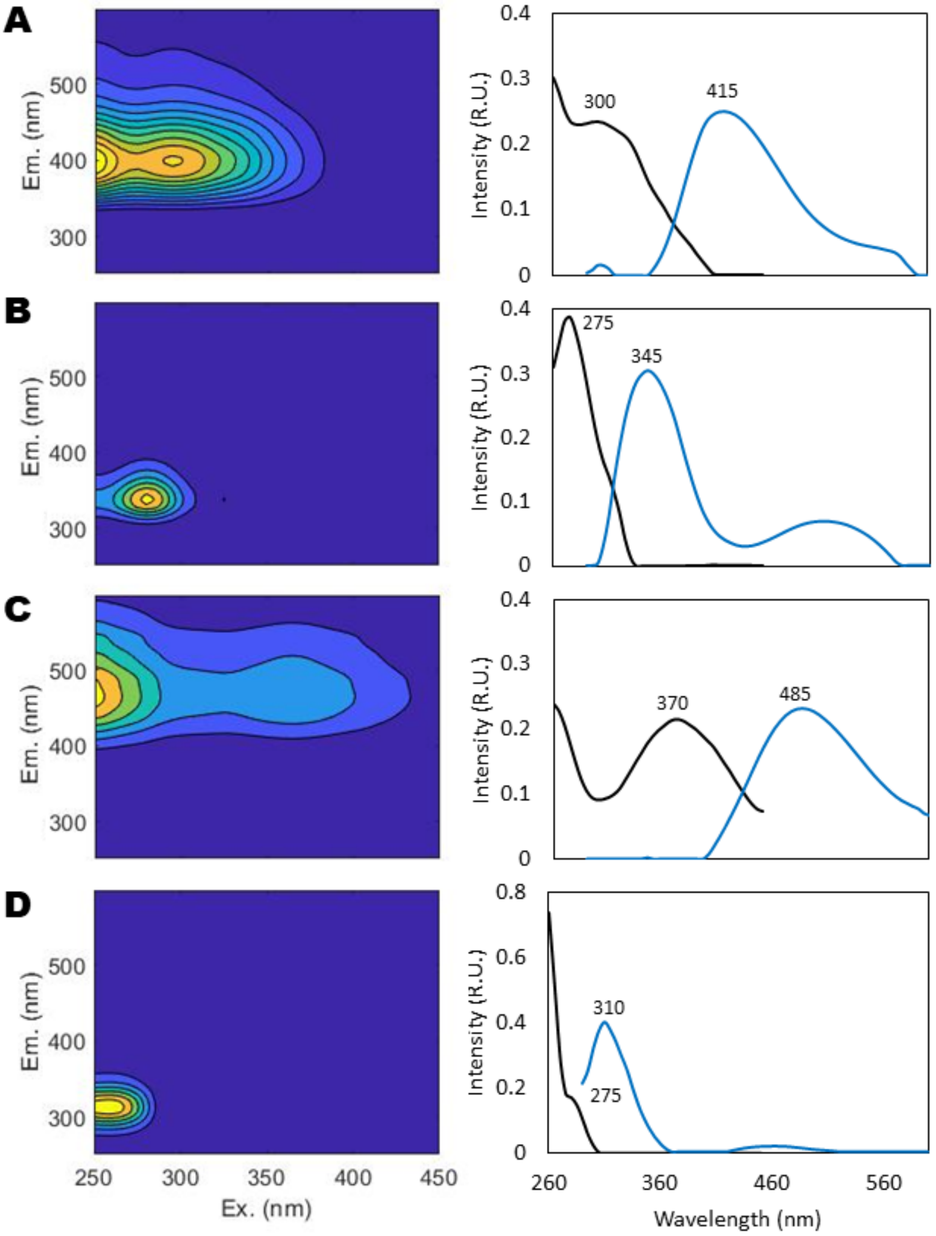


**Fig. S.I. 1: Fluorescence signatures of the four PARAFAC components identified in the dataset:** (**A**) C1 marine humic-like, (**B**) C2 tryptophan-like, (**C**) C3 terrestrial fulvic-like, (**D**) C4 Tyrosine- like. Contour plots of components C1 - C4 are ordered by decreasing percent explained, with emission wavelength on the y-axis, excitation on the x-axis, and shading representing the relative intensity of emission. Corresponding line plots to the right of each contour plot represent the excitation (black curve) and emission (blue curve) spectra of each component. The x-axis in the line plots is excitation or emission wavelength, with relative intensity on the y-axis.

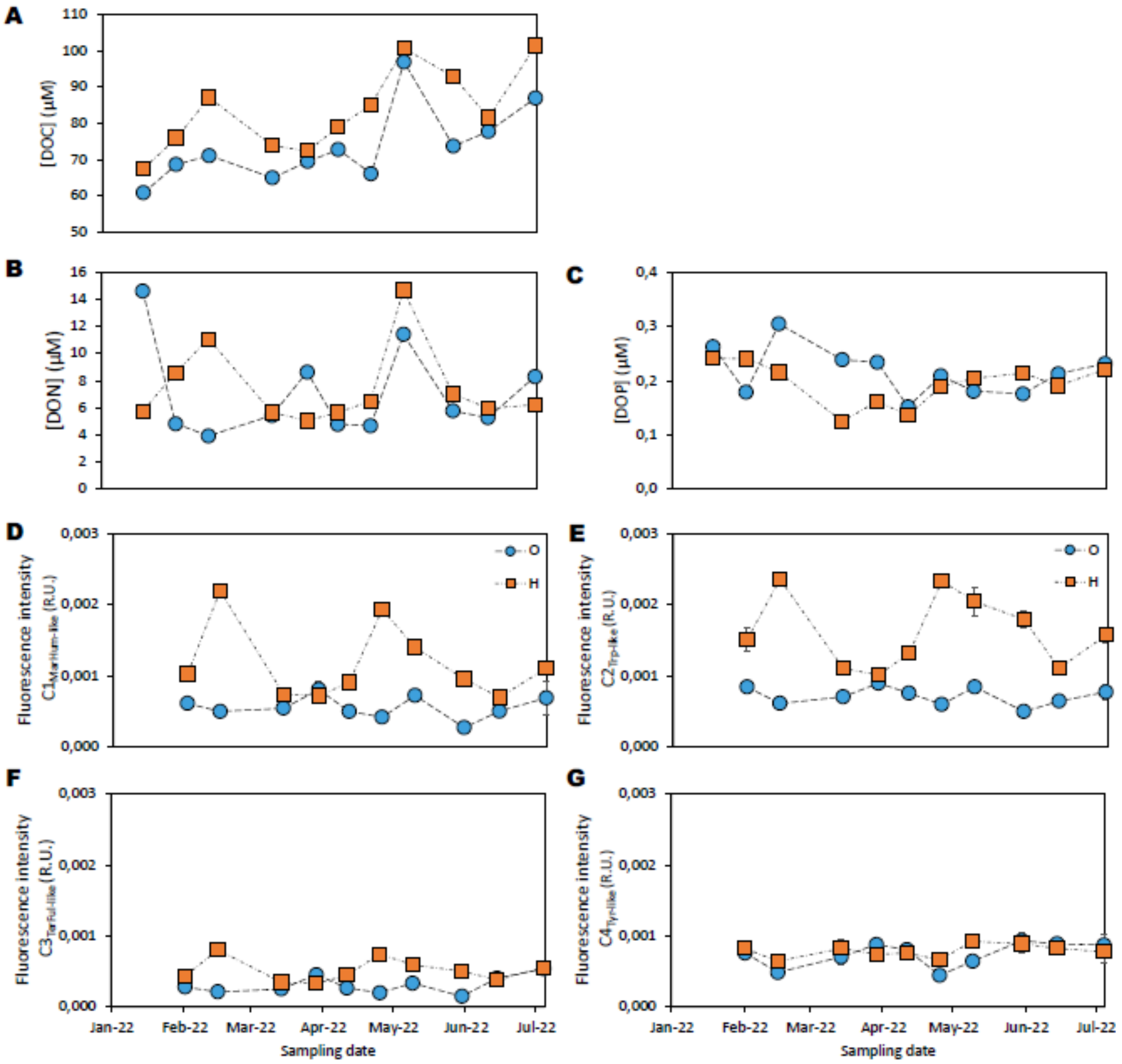


**Fig. S.I. 2: Comparison of the temporal dynamics of abiotic parameters between harbour area (□) and open area (○).** (A) Dissolved Organic Carbon (DOC), (B) Dissolved Organic Nitrogen (DON), (C) Dissolved Organic Phosphorus (DOP), and Fluorescence intensity of (D) $C_{1MarHum\text{-}like}$, (E) $C_{2Trp\text{-}like}$, (F) $C_{3TerFul\text{-}like}$ and (G) $C_{4Tyr\text{-}like}$ identified by PARAFAC analysis. Error bars denote standard error of triplicate analyses.

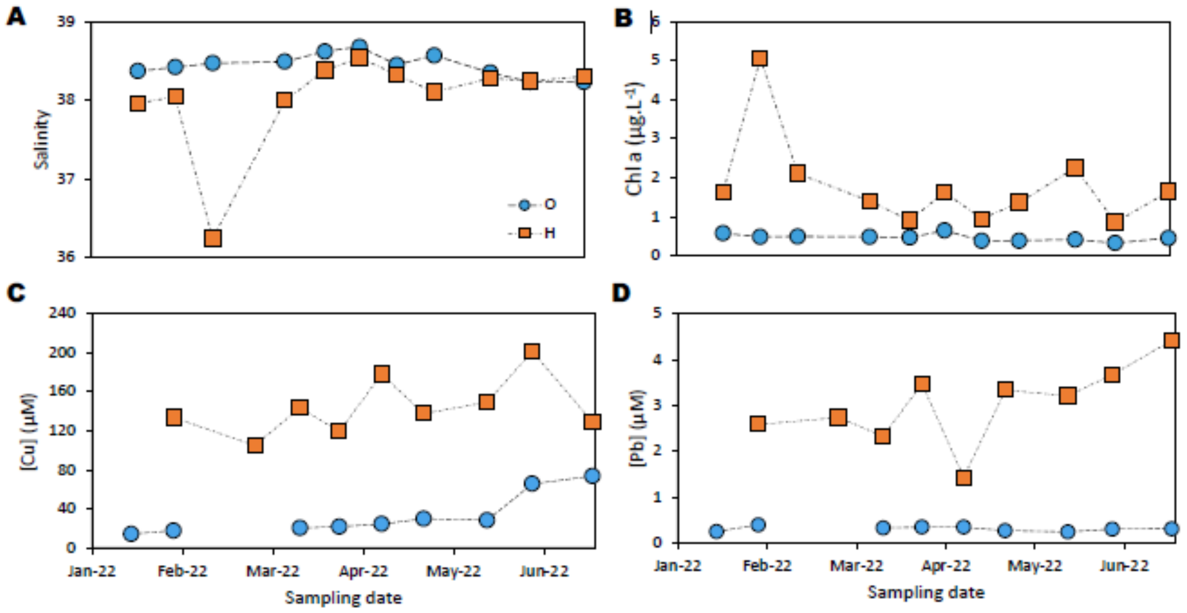


**Fig. S.I. 3: Comparison of the temporal dynamics of abiotic parameters between harbour area (□) and open area (○).** (A) Salinity, (B) Chlorophyll a, (C) Copper (Cu) concentration, (D) Lead (Pb) concentration.

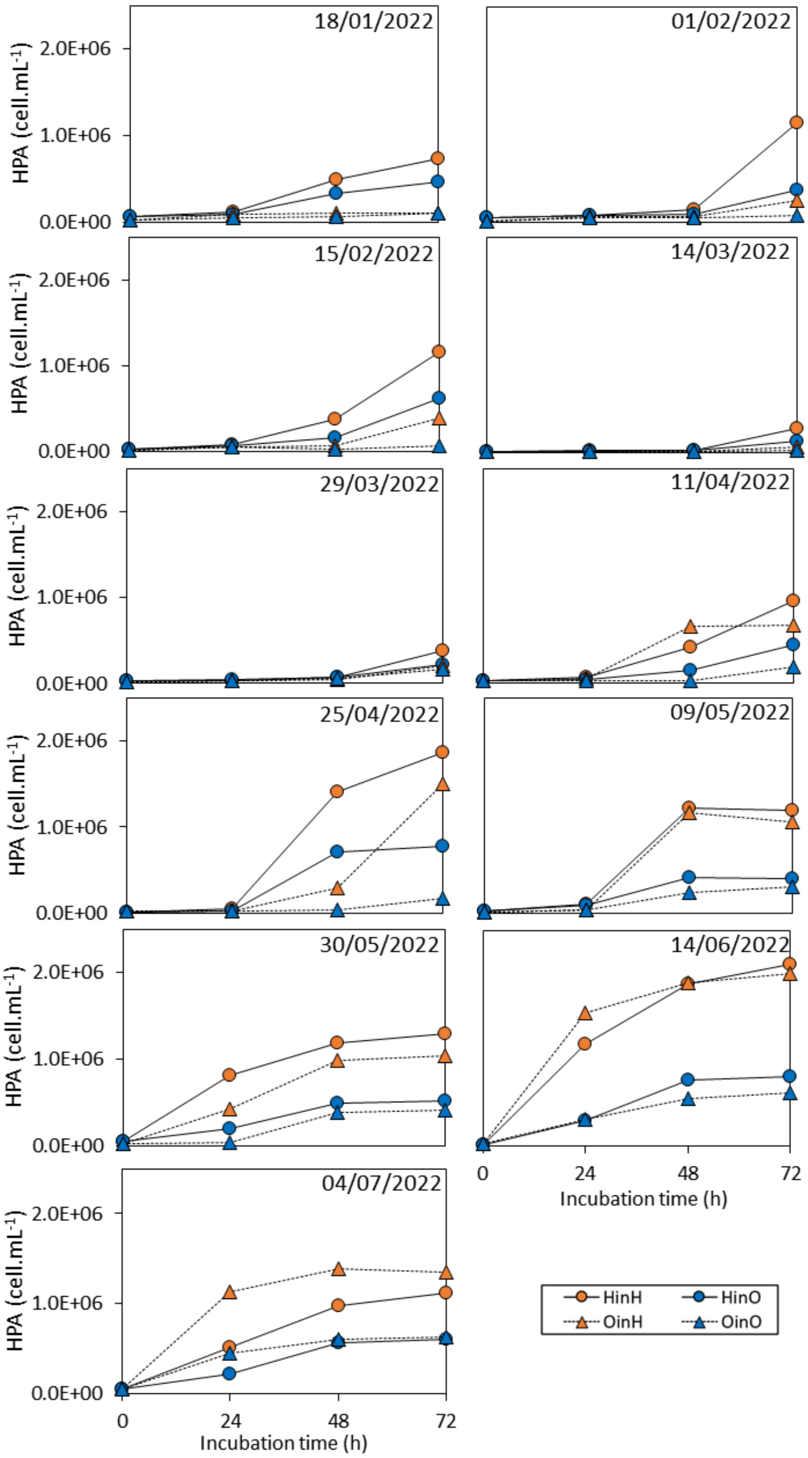


**Fig. S.I. 4: Heterotrophic prokaryote abundances (HPA) as a function of DOM and community origin for each campaign**. HP origin is specified by point shapes. Circles: Harbour HP. Triangles: Open area HP. DOM origin is specified by point colours. Orange: Harbour DOM. Blue: Open area DOM. Error bars represent the standard deviation between experimental triplicates.

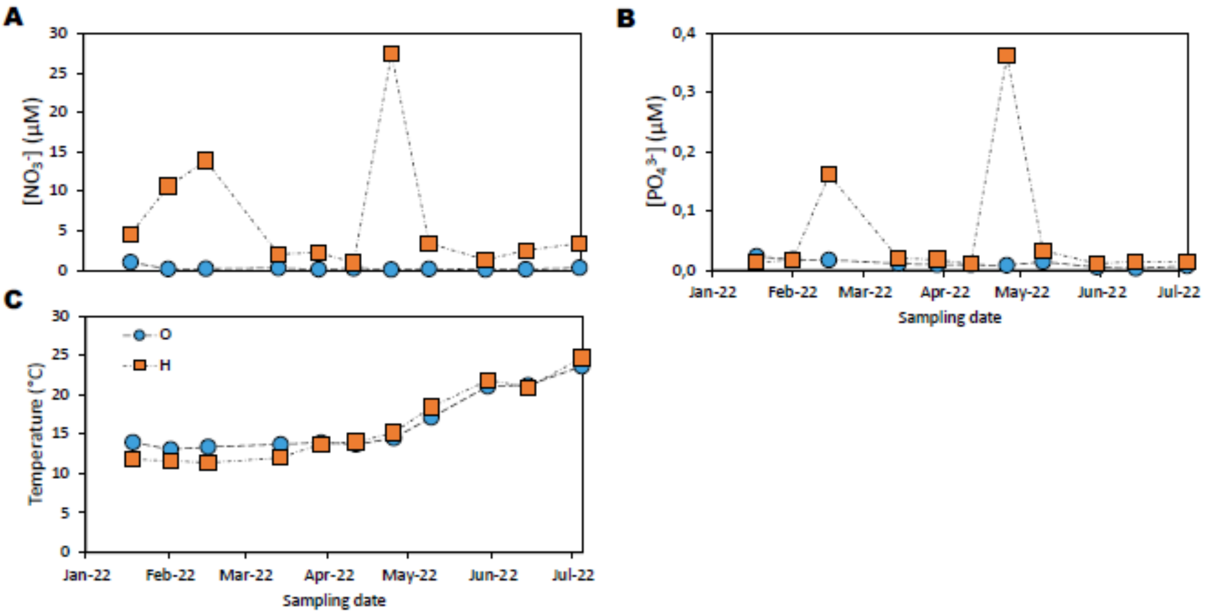


**Fig. S.I. 5: Comparison of the temporal dynamics of abiotic parameters between harbour area (□) and open area (○).** (A) Nitrate ($NO_3^-$), (B) Phosphate ($PO_4^{3-}$), (C) Temperature.
